# Advanced neural activity mapping in brain organoids via field potential imaging with ultra-high-density CMOS microelectrodes

**DOI:** 10.1101/2025.05.24.655914

**Authors:** Remi Yokoi, Naoki Matsuda, Yuto Ishibashi, Ikuro Suzuki

**Author notes:** **Correspondence:** Ikuro Suzuki.

## Abstract

Human iPSC-derived brain organoids and assembloids have emerged as promising in vitro models for recapitulating human brain development, neurological disorders, and drug responses. However, detailed analysis of their electrophysiological properties requires advanced measurement techniques. Here, we present a novel analytical approach utilizing ultra-high-density (UHD) CMOS microelectrode arrays (MEAs) containing 236,880 electrodes (10.52 μm × 10.52 μm each) distributed over a broad sensing area of 32.45 mm^2^ for field potential imaging (FPI) of brain organoids. Neuronal activity was recorded simultaneously from over 46,000 electrodes interfaced with brain organoids, allowing for the identification of single-cell firing events and the assessment of neuronal network connectivity based on individual spikes. In midbrain organoids, administration of L-DOPA revealed both excitatory and inhibitory cellular responses, with a dose-dependent increase in the proportion of excitatory responses, suggesting enhanced network connectivity. Capitalizing on the spatial and temporal resolution of UHD-CMOS-MEAs, we introduced new endpoints for network activity: propagation velocity and propagation area. In cortical organoids, application of the GABA_A_ receptor antagonist picrotoxin led to increased propagation velocity, whereas the NMDA receptor antagonist MK-801 resulted in a broad reduction of propagation area, along with localized increases. As FPI enables direct recording of electrical potential waveforms, frequency-domain analyses were also conducted. Spontaneous activity in cortical organoids exhibited region-specific frequency distributions, with gamma-band activity displaying distinct patterns compared to other frequency bands. Additionally, in midbrain–striatal assembloids, electrophysiological activity was observed in both regions. Connectivity analysis showed that treatment with 4-aminopyridine enhanced inter-organoid connection strength. This large-scale, single-cell-resolved recording approach using UHD-CMOS-MEAs facilitates comprehensive analysis of network connectivity, propagation velocity and propagation area, and frequency characteristics. It represents a powerful platform for advancing our understanding of the electrophysiological functions of brain organoids and assembloids, and holds significant potential for drug screening and disease modeling in human neuroscience research.

## 1 Introduction

The human brain exhibits remarkably complex structures and functions, and the diverse dynamics of neural activity are intricately involved in development, cognition, and disease. Brain organoids have recently attracted attention as in vitro models that enable the reconstruction and analysis of these brain functions. They are three-dimensional cultured tissues derived from pluripotent stem cells that self-organize into neural structures.(Lancaster et al., 2013; Lancaster and Knoblich, 2014; Giandomenico et al., 2021) To date, region-specific brain organoids mimicking the cerebral cortex (Lancaster et al., 2013), striatum (Miura et al., 2020), midbrain (Jo et al., 2016), and hippocampus (Sakaguchi et al., 2015) have been developed, allowing for the recapitulation of developmental processes and functional assessment of neural networks. In the context of compound screening, brain organoids are expected to serve as novel platforms that enable more precise prediction and evaluation of drug efficacy and toxicity than conventional two-dimensional cultures or animal models. Furthermore, because brain organoids can be generated from patient-derived induced pluripotent stem cells (iPSCs), they provide an opportunity to model human-specific disease phenotypes that are difficult to reproduce in animal systems. Indeed, various disease-related features have been reported using brain organoids, including amyloid-β and tau pathology in Alzheimer’s disease (Raja et al., 2016; Gonzalez et al., 2018; Yan et al., 2018), overproduction of GABAergic neurons in autism spectrum disorder (Mariani et al., 2015), reduced neural and increased neurovascular cell populations in schizophrenia (Notaras et al., 2022), and disease-specific frequency characteristics and responses to contraindicated drugs in Dravet syndrome (Yokoi et al., 2023).

To construct more structurally complex in vitro models, increasing efforts have been directed toward generating assembloids by fusing organoids derived from different brain regions. Various types of assembloids have been developed to date, including cortical–striatal (Miura et al., 2020), midbrain– striatal (Ozgun et al., 2024), and midbrain–striatal–cortical assembloids (Reumann et al., 2023). More recently, in 2025, a model of the ascending neural sensory pathway was established by connecting organoids representing the somatosensory system, spinal cord, thalamus, and cortex(Kim et al., 2025).

As a functional application of brain organoids, their potential use in bio-computing systems has attracted growing interest. In 2022, the concept of “Organoid Intelligence” was proposed as an emerging interdisciplinary field that aims to harness the self-organizing capabilities of brain organoids to perform learning and memory functions within bio-computing platforms (Smirnova et al., 2023).

To achieve these goals, it is essential to establish detailed methods for evaluating the electrical activity of brain organoids. Currently, electrophysiological functional assessments of organoids employ techniques such as patch-clamp recordings, calcium imaging, and microelectrode array (MEA) recordings. MEA-based approaches allow for long-term, noninvasive monitoring of the electrical activity of brain organoids and assembloids, enabling the observation of time-dependent changes in network dynamics and the evaluation of pharmacological responses (Tasnim and Liu, 2022; Lv et al., 2023; Pan et al., 2024). Moreover, analysis of low-frequency signals (below 500 Hz) has been shown to be effective for assessing pharmacological responses in brain organoids (Yokoi et al., 2021). In addition, disease-specific frequency characteristics and responses to contraindicated compounds have also been successfully detected (Yokoi et al., 2023). Recently, the development of high-density (HD) CMOS-based MEAs utilizing complementary metal-oxide semiconductor (CMOS) technology has further improved the precision of neuronal network activity evaluation. Recordings from brain organoids using HD CMOS MEAs have enabled the quantification of dynamic single-cell firing rates and the calculation of propagation velocities (Schröter et al., 2022; Sharf et al., 2022). However, these evaluations have generally been limited to small recording areas and a restricted number of electrodes, falling short of capturing activity across the entire organoid.

In this study, we present an analytical approach for detailed evaluation of electrical activity in brain organoids using field potential imaging (FPI) with an ultra-high-density (UHD) CMOS microelectrode array (MEA) comprising 236,880 microelectrodes (10.52 μm × 10.52 μm) across a wide sensing area of 32.45 mm^2^ (Suzuki et al., 2023). Specifically, we conducted (1) single-cell spike identification across the entire organoid and evaluation of neuronal network connectivity based on spike activity; (2) calculation of propagation velocity and propagation area as novel endpoints of network activity; and (3) spatial analysis of frequency-specific signal characteristics. Furthermore, we applied the UHD-CMOS MEA to midbrain–striatal assembloids to record electrical activity from both tissues and analyze inter-regional connectivity.

## 2 Materials and Methods

### 2.1 Culture of Human iPS Cells

Human induced pluripotent stem cells (hiPSCs) derived from a healthy donor (201B7 line, obtained from RIKEN) were cultured in 6-well plates pre-coated with Vitronectin (07180, STEMCELL Technologies) using mTeSR™ Plus medium (100-0276, STEMCELL Technologies). The culture medium was partially replaced (50%) every 3 to 4 days. Once the cells reached confluence, they were dissociated using Gentle Cell Dissociation Reagent (07174, STEMCELL Technologies) and subsequently used for organoid generation.

### 2.2 Culture of Midbrain Organoids

Midbrain organoids used in this study were pre-generated and supplied by STEMCELL Technologies. After acquisition, the organoids were cultured for four months using the STEMdiff™ Neural Organoid Maintenance Kit (100-0120, STEMCELL Technologies). The culture medium was then replaced with BrainPhys™ Neuronal Medium (05792, STEMCELL Technologies), and complete medium changes were carried out every 3 to 4 days to maintain the organoids.

### 2.3 Culture of Cerebral Organoids

Cortical organoids were generated using the STEMdiff™ Cerebral Organoid Kit (08570, STEMCELL Technologies). Human iPS cells were seeded at a density of 9.0 × 10^3^ cells per well, using EB seeding medium. On days 2 and 4 of culture, 100 µL of EB formation medium was added to each well. On day 5, the medium was replaced with induction medium, and the cultures were incubated for 2 days. On day 7, organoids were embedded in Matrigel (354277, Corning) and incubated in expansion medium for 3 days. From day 10 onward, the expansion medium was replaced with maturation medium, and the organoids were cultured on an orbital shaker (COSH6, AS ONE Corporation). Organoids were maintained in maturation medium for 3 months, with medium changes carried out every 3 to 4 days. After 3 months, the culture medium was switched to BrainPhys™ Neuronal Medium (05792, STEMCELL Technologies) for long-term maintenance. Details of the media components and kit reagents are provided in Supplementary Table 1.

### 2.4 Culture of Midbrain–Striatal Assembloids

Midbrain–striatal assembloids were generated using the STEMdiff™ Midbrain Organoid Differentiation Kit (100-1096, STEMCELL Technologies) and the STEMdiff™ Dorsal Forebrain Organoid Differentiation Kit (08620, STEMCELL Technologies). To induce striatal differentiation, organoids were cultured in medium supplemented with Activin A, IWP-2, and SR11237, following the manufacturer’s recommended conditions. Human iPSCs were seeded at a density of 3 × 10^6^ cells per well in Organoid Formation Medium supplemented with 10 µM Y-27632, using AggreWell™800 24-well plates (34811, STEMCELL Technologies) pretreated with AggreWell Rinsing Solution (07010, STEMCELL Technologies). From days 2 to 5, half-medium changes were performed. On day 6, the culture medium was replaced with Midbrain Organoid Expansion Medium for midbrain organoids and Striatal Organoid Expansion Medium for striatal organoids. Full medium changes were performed every other day until day 25. On day 25, the respective Organoid Differentiation Media were introduced, and the same schedule of complete medium changes continued until day 43. From day 43 onward, cultures were maintained in Organoid Maintenance Medium. On day 50 (midbrain) and day 60 (striatal), one organoid of each type was placed together in a well of a 24-well plate and co-cultured on an orbital shaker (COSH6, AS ONE Corporation) to form midbrain–striatal assembloids. These assembloids were maintained in Maintenance Medium for an additional four months, with medium changes every 3 to 4 days. Details of the media components and kit reagents are provided in Supplementary Table 2.

### 2.5 Immunocytochemistry

Cultured brain organoids were fixed in 4% paraformaldehyde (PFA) in phosphate-buffered saline (PBS). After fixation, the organoids were embedded in Optimal Cutting Temperature (OCT) compound (45833, Sakura Finetek Japan), and 10 µm-thick cryosections were prepared using a cryostat (CM1950, Leica). The sections were first permeabilized with 0.2% Triton X-100 for 10 min and then incubated in pre-blocking buffer (PBS containing 0.05% Triton X-100 and 5% goat serum) at 4 °C for 60 min. Primary antibody incubation was carried out in pre-blocking buffer at 4 °C for 12 hours using anti-MAP2 antibody (ab5392, Abcam) and anti-tyrosine hydroxylase (TH) antibody (ab137869, Abcam). For fluorescence detection, Alexa Fluor 488-conjugated anti-rabbit IgG (A11040, Invitrogen) and Alexa Fluor 546-conjugated anti-chicken IgG (A11008, Invitrogen) were diluted 1:1000 in pre-blocking buffer and incubated at room temperature for 1 hour. Nuclear staining was performed with 1 µg/mL Hoechst 33258 (343-07961, Dojindo) for 1 hour at room temperature. Fluorescently labeled organoid sections were imaged using a confocal laser scanning microscope (AX R, Nikon), and image analysis was performed using ImageJ software (NIH).

### 2.6 Ultra-high-density CMOS MEA Recording

In this study, the electrical activity of brain organoids was recorded using the field potential imaging (FPI) technique with an ultra-high-density CMOS-based microelectrode array (UHD-CMOS MEA) system (Suzuki et al., 2023). The MEA chip comprised 236,880 electrodes, each measuring 10.52 µm × 10.52 µm (110.67 µm^2^), with a total active recording area of 32.45 mm^2^. Cerebral organoids, midbrain organoids, and midbrain–striatal assembloids were individually mounted on the CMOS-MEA system (Sony Semiconductor Solutions), and recordings were conducted under controlled conditions (5% CO_2_, 37 °C). Electrical signals were acquired at a sampling rate of either 2 kHz or 10 kHz. Baseline drift was removed using least-squares detrending applied to each 500-sample segment, followed by offset correction to stabilize the signal.

### 2.7 Pharmacological Assays

Pharmacological experiments were conducted to evaluate the responsiveness of brain organoids to neurotransmitter-related agents. In 4-month-old midbrain organoids, 3-(3,4-dihydroxyphenyl)-L-alanine (L-DOPA; D0600, Tokyo Chemical Industry), a dopamine precursor, was applied, and spontaneous electrical activity was recorded for 2 min at each concentration. In 5-month-old cerebral organoids, the GABA_A_ receptor antagonist picrotoxin (PTX; 168-17961, FUJIFILM Wako) and the NMDA receptor antagonist MK-801 (M107-50MG, Sigma) were administered separately, and spontaneous activity was recorded for 1 min at each concentration. Additionally, to assess changes in inter-regional connectivity within midbrain–striatal assembloids, 4-aminopyridine (4-AP; 016-02781, FUJIFILM Wako) was applied to 4-month-old assembloids, and spontaneous activity was recorded for 2 min at each concentration following 4-AP application.

### 2.8 Spike Detection and Soma identification

Spike detection was performed on voltage signals processed with a 100 Hz high-pass filter, using a threshold of ±5.0σ, where σ denotes the standard deviation of baseline noise during a quiescent period. Events exceeding this threshold were defined as spikes. For each detected spike, a waveform segment comprising 20 samples before and after the spike time point (41 samples in total) was extracted. Waveforms with a standard deviation below 13 were considered noise and excluded from further analysis. Soma identification was then performed based on spike timing correlations between electrode pairs. For all electrode pairs in which spikes were detected, inter-spike intervals (ISIs) were calculated. If the ISI between spikes on two electrodes was less than 10 ms, it was counted as a same-count event. Electrode pairs with ≥5 same-counts that also constituted ≥15% of all possible spike pairs were designated as putative somatic electrode pairs. If the distance between electrodes in a putative pair was less than 50 µm, they were considered to originate from the same neuronal soma, and the location was identified as a somatic candidate site. In the case of midbrain organoids, spike events occurring during network bursts were excluded from soma identification to minimize false positives.

### 2.9 Z-score Analysis

For each neuronal pair, a synchronous spike was defined as an event in which a spike from one cell was followed by a spike from the other cell within 100 ms. The total number of synchronous spikes was counted for each pair. To assess statistical significance, 100 surrogate spike trains were generated by randomly shuffling the original spike time sequences while preserving each cell’s inter-spike interval (ISI) distribution. For each surrogate dataset, the number of synchronous spikes was calculated using the same criteria. A z-score was calculated using the mean and standard deviation of the synchronous spike counts across the surrogate datasets, based on the synchronous spike count in the actual data. A z-score ≥ 3 was considered to indicate significant spike synchrony between the two cells, and such pairs were defined as having a functional connection. Based on the results of the z-score analysis, the number of significantly connected partner cells was counted for each neuron to determine its connection degree.

### 2.10 Propagation Analysis

Oscillatory activity was detected in voltage waveforms recorded using the UHD-CMOS MEA. Electrodes that exhibited signal amplitudes ≥30 µV during these periods were defined as propagating electrodes, while all others were treated as noise electrodes and excluded from the analysis. For each propagating electrode, the peak time—defined as the interval from the onset of oscillation (t = 0) to the point of maximum amplitude—was calculated. A time-series dataset was then constructed by counting the number of electrodes that reached their peak within each time bin. The propagation velocity of network activity was estimated by computing the temporal derivative of this time series. Propagation velocity was calculated for each oscillatory event detected in individual traces, and the average of these values was reported as the final result.

### 2.11 Frequency Band Power Analysis of Oscillatory Waveforms

Detected oscillatory waveforms were processed using zero-phase filtering with a finite impulse response (FIR) bandpass filter to isolate signals within distinct frequency bands. The potential amplitude for each frequency band was calculated from the filtered waveforms. The five frequency bands analyzed were defined as follows: delta (0.5–3 Hz), theta (4–7 Hz), alpha (8–11 Hz), beta (12– 29 Hz), and gamma (30–100 Hz). For each band, power values were standardized using the mean and standard deviation across all electrodes. The resulting normalized data were subjected to unsupervised clustering using the k-means algorithm implemented in MATLAB (MathWorks) to classify patterns of frequency-specific potential dynamics.

### 2.12 Analysis of Connection Strength in Midbrain–Striatal Assembloids

For each neuronal pair, a synchronous spike was defined as a spike from one cell followed by a spike from the other cell within 100 ms. The number of synchronous spikes was counted for each pair. To minimize the influence of differences in firing rates between cells, the synchronous spike count was normalized by the mean spike count of the respective cell pair. The resulting value was defined as the connection strength. Connection strength was calculated separately for three groups: striatal organoids, midbrain organoids, and midbrain–striatal assembloids.

## 3 Results

### 3.1 Single-Cell Firing Rate and Intercellular Connectivity Analysis of Midbrain Organoids Following L-DOPA Administration

Immunofluorescence staining of midbrain organoids derived from human iPSCs and cultured for 4 months confirmed the expression of MAP2, a marker for mature neurons, and TH, a marker for dopaminergic neurons (Figure. 1A). In addition, non-TH-positive neurons were present. The midbrain organoid was mounted on a UHD-CMOS MEA (Figure. 1B), and spontaneous electrical activity was recorded across the entire organoid, resulting in the identification of 404 individual cells. A raster plot and histogram of the spontaneous spike activity over a 2-minute period from these identified cells are shown in Figure. 1C. Network bursts, in which almost all cells spiked simultaneously, were observed (indicated by peaks in the histogram and vertical black lines in the raster plot in Figure. 1C), suggesting the formation of functional neuronal circuits within the midbrain organoid. Figure 1D presents a spatial map of total spike counts over 2 minutes for the 404 identified cells, with dot size representing the number of spikes. Cells whose spike count increased to ≥150% following administration of L-DOPA (30 µM) compared to before treatment are shown in red, while those with a decrease to ≤50% are shown in blue. The proportion of cells with increased activity was 19.3% at 0.3 µM, 39.4% at 3 µM, and 59.7% at 30 µM. Conversely, the proportion of cells with decreased activity was 10.1%, 2.23%, and 0.495%, respectively, while the proportions of non-responsive cells were 70.5%, 58.4%, and 39.9%, respectively (Figure. 1E). The single-cell resolution afforded by the UHD-CMOS MEA enabled the detection not only of increased spike activity following L-DOPA treatment but also of cells that were non-responsive or showed suppressed activity at lower doses. To evaluate network activity in the midbrain organoid, z-scores were calculated for cell pairs that spiked within 100 ms. In Figure. 2A, black lines indicate cell pairs with a z-score ≥ 3 before L-DOPA administration, while blue lines indicate pairs whose z-scores increased after treatment. L-DOPA administration led to a dose-dependent increase in connection strength (Figure. 2B). The distribution of z-scores also shifted to the right in accordance with L-DOPA concentration (Figure. 2C), suggesting enhanced synchrony within the network. To quantitatively assess changes in connection strength, cell pairs with z-scores ≥ 3 were defined as connected, and the average number of connected cells per cell was calculated. The number of connections per cell was 333.2 ± 84.6 at baseline, 335.8 ± 79.2 at 0.3 µM, 380.5 ± 42.8 at 3 µM, and 382.7 ± 45.2 at 30 µM, with significant increases observed at 3 µM and 30 µM compared to baseline (Figure. 2D). These findings indicate that the UHD-CMOS MEA enables both single-cell spike analysis and intercellular connectivity evaluation in brain organoids.

**Figure 1.**
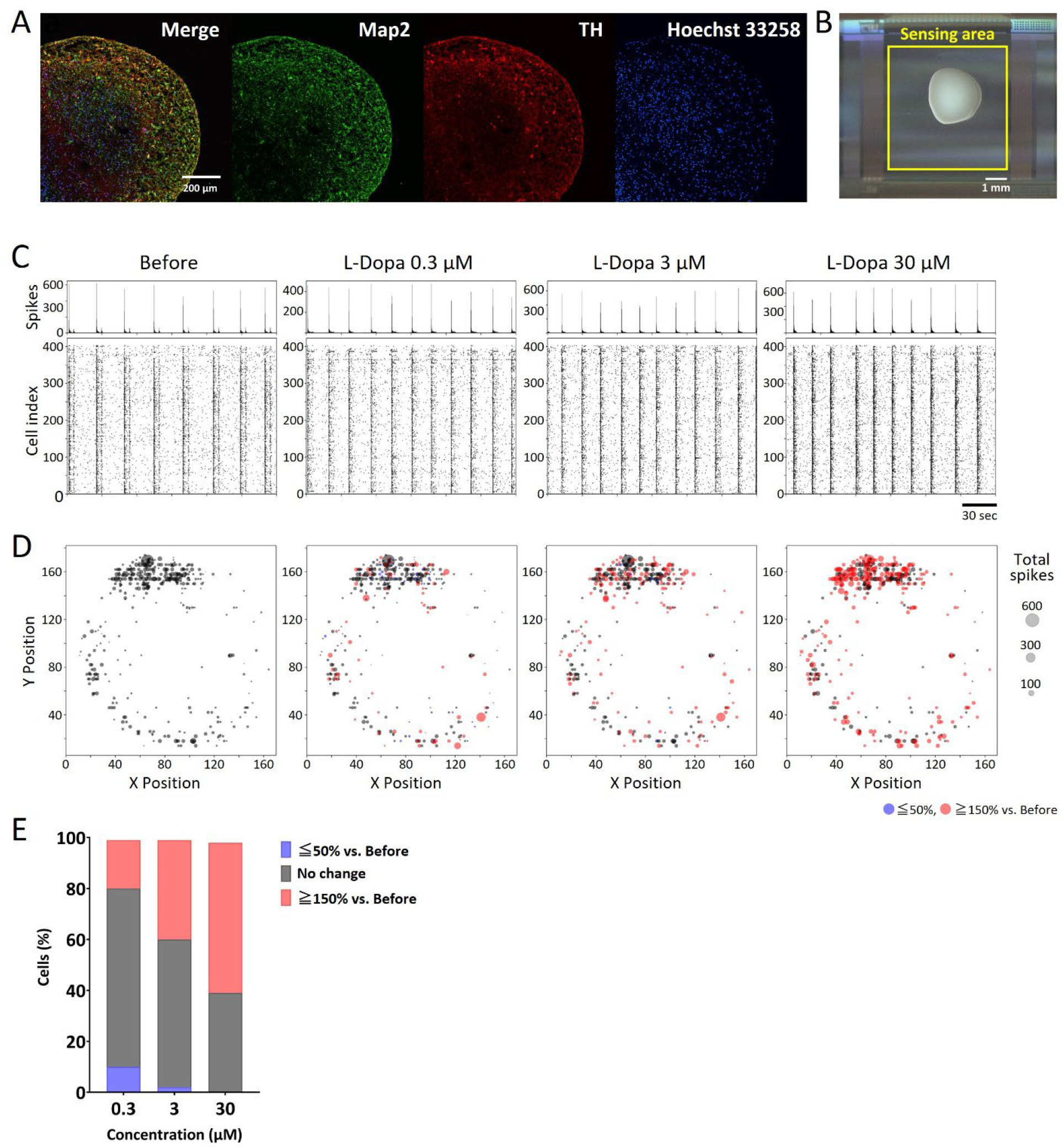
Single-cell spike analysis of midbrain organoids. (A) Immunofluorescence image of a human midbrain organoid cultured for 4 months. Green: MAP2; red: TH; blue: Hoechst 33258. Scale bar = 200 µm. (B) Midbrain organoid mounted on a UHD-CMOS MEA. The sensing area is indicated by a yellow square. Scale bar = 1 mm. (C) Raster plot and histogram of spontaneous neuronal spikes over a 2-minute period before and after L-DOPA administration. Scale bar = 30 sec. (D) Spatial map of spontaneous spikes over a 2-minute period from 404 detected cells. Each dot represents a cell body, and dot size reflects spike frequency. Red: ≥150% vs. before; blue: ≤50% vs. before. (E) Change in spike frequency following L-DOPA administration. The proportion of cells showing altered activity relative to before administration is indicated. Red: ≥150% vs. before; blue: ≤50% vs. before; black: no change.

**Figure 2.**
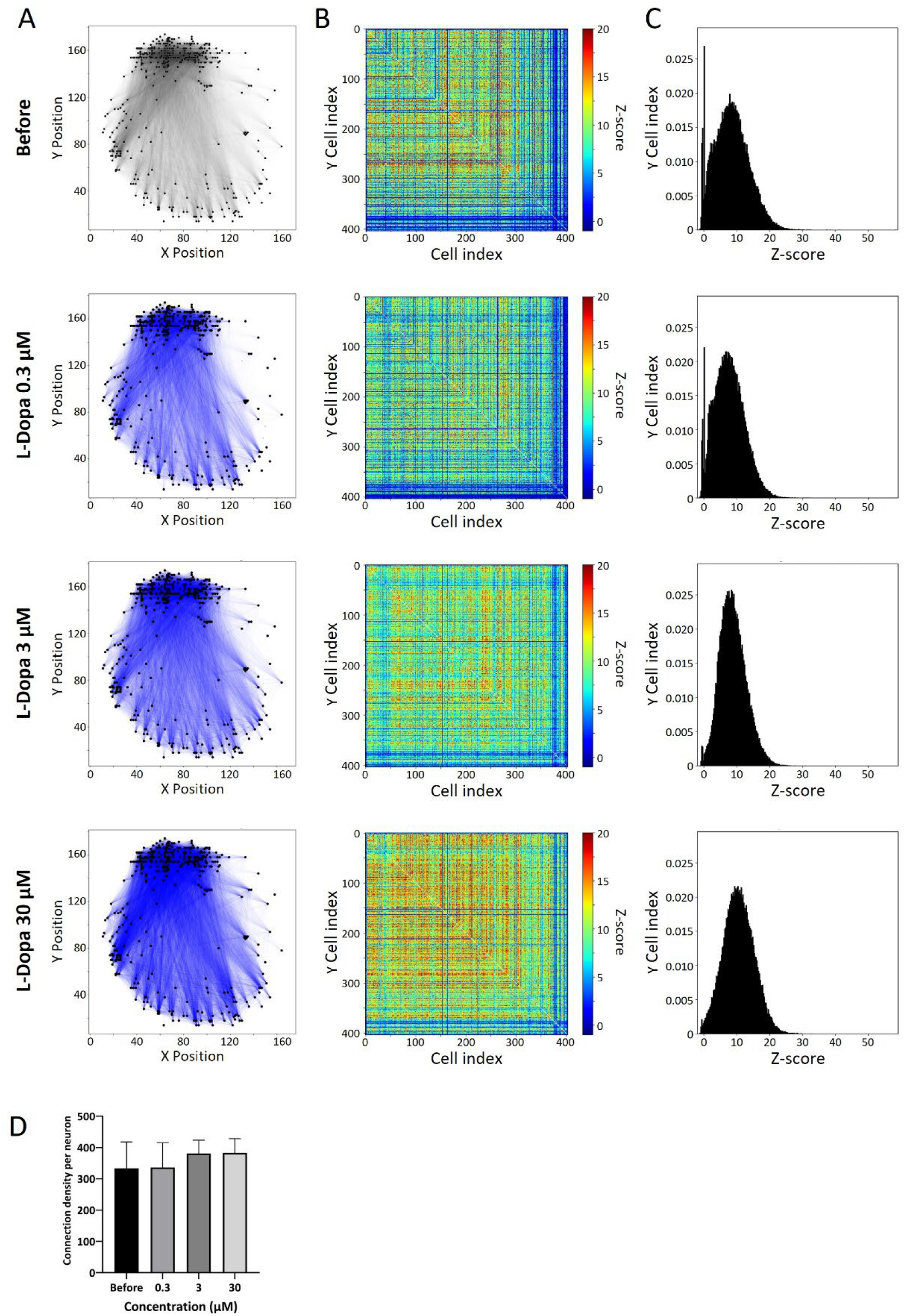
Neural network analysis of midbrain organoids. Changes in connection strength between cell pairs before and after L-DOPA administration. To evaluate network activity, z-scores were calculated for cell pairs that spiked within 100 ms. (A) Cell connection map. Black lines indicate cell pairs with a z-score ≥ 3; blue lines indicate pairs whose z-score increased after L-DOPA treatment. (B) Heatmap of z-scores for 404 detected cells. (C) Distribution of z-scores. (D) Change in the number of connected cells per individual cell.

### 3.2 Analysis of Propagation Velocity and Propagation Area in Cortical Organoids

The ultra-high-density (UHD) CMOS MEA, with its high temporal and spatial resolution, enabled the derivation of novel endpoints for evaluating network activity in brain organoids—specifically, propagation velocity and propagation area. To assess propagation velocity, the GABA_A receptor antagonist picrotoxin (PTX) was applied to 5-month-old cerebral organoids. Figure 3A shows representative propagation waveforms recorded during spontaneous oscillations, with the corresponding electrode positions indicated by red arrows in the top-left panel of Figure. 3B (Before). Peak timing maps of oscillatory waveforms across electrodes, recorded before and after PTX administration, are presented in Figure. 3B, illustrating that neural activity propagated throughout the entire organoid. Propagation velocity was calculated by differentiating the number of electrodes reaching peak voltage per unit time (Figure. 3C). The baseline propagation velocity was 0.0166 ± 0.000715 mm^2^/ms, which significantly increased to 0.0202 ± 0.000523, 0.0193 ± 0.000394, and 0.0196 ± 0.000257 mm^2^/ms following administration of 0.1, 1, and 10 µM PTX, respectively (Figure. 3C). Propagation area was assessed before and after treatment with the NMDA receptor antagonist MK-801. In the maps shown in Figure. 3D, electrodes exhibiting propagation activity are indicated in black. At 0.1 µM MK-801, the propagation area increased from 0.851 mm^2^ to 1.05 mm^2^, although the change was not statistically significant (Figure. 3E). By contrast, at 1 µM MK-801, the propagation area decreased markedly across most of the organoid, with a localized increase observed only in the lower right region (Figure. 3D). At the whole-organoid level, synaptic transmission blockade resulted in a significant reduction of the propagation area to 0.384 mm^2^ (Figure. 3E). These findings indicate that the analysis of propagation velocity and propagation area using UHD-CMOS MEAs provides an effective approach for evaluating compound-induced responses related to synaptic transmission in brain organoids.

**Figure 3.**
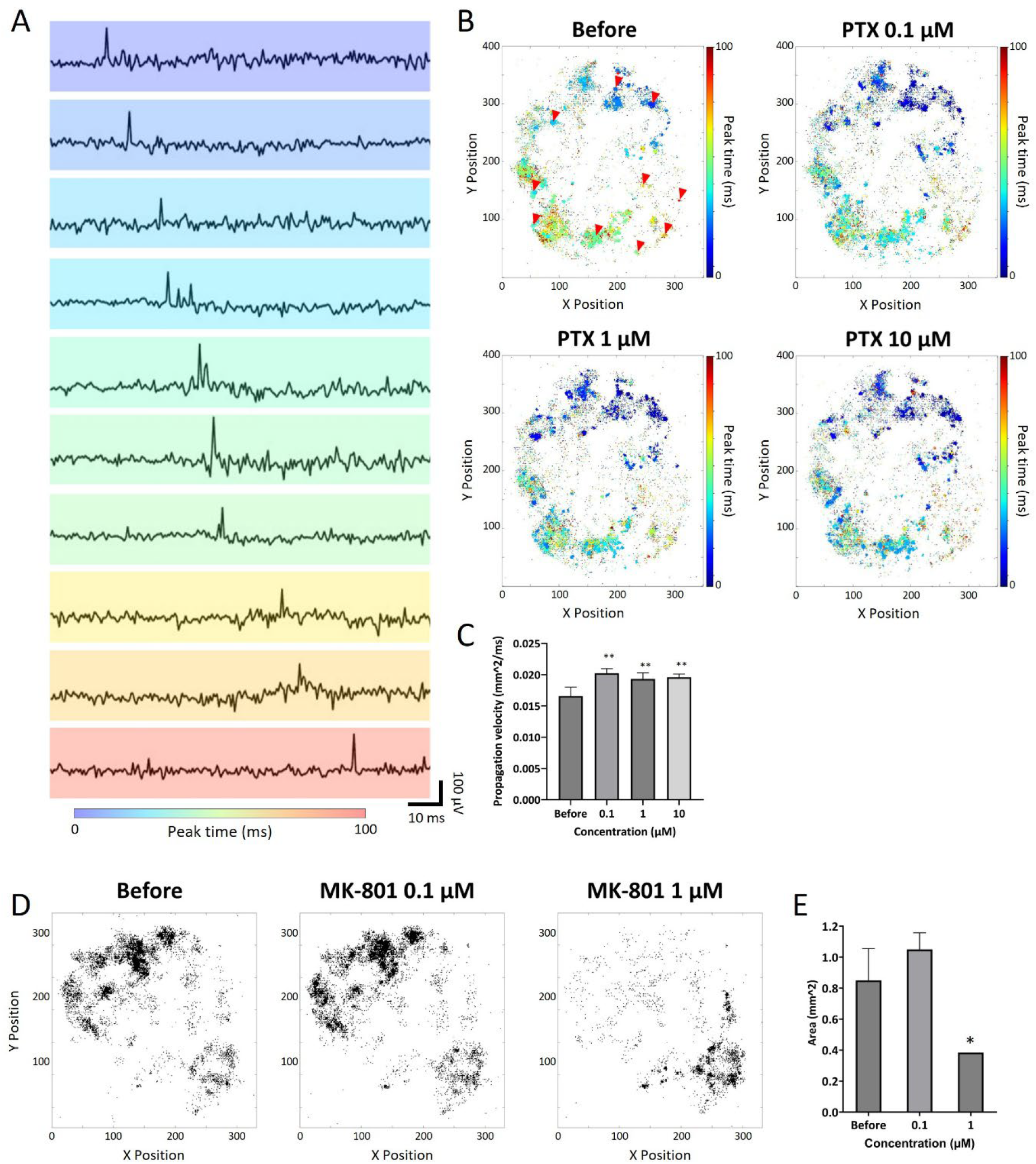
Propagation analysis of cortical organoids. (A) Spontaneous activity propagation waveform color-coded by the timing of voltage peaks. The electrode used for pickup is indicated by a red arrow in (B) Before. (B) Delay maps of oscillation peak timing before and after picrotoxin (PTX) administration. (C) Propagation velocity before and after PTX administration. The average propagation velocity was calculated per oscillation event (before: n = 4; 0.1 µM: n = 2; 1 µM: n = 6; 10 µM: n = 4). Data were analyzed using one-way ANOVA followed by post hoc Dunnett’s test (**p < 0.01 vs. before). (D) Electrode maps showing propagation before and after MK-801 administration; black dots indicate propagation electrodes. (E) Propagation area before and after MK-801 administration. The propagation area was calculated per oscillation event (before: n = 6; 0.1 µM: n = 6; 1 µM: n = 1). Data were analyzed using one-way ANOVA followed by post hoc Dunnett’s test (*p < 0.05 vs. before).

### 3.3 Frequency Characteristics of Cerebral Organoids

Field potential imaging (FPI) using the UHD-CMOS MEA provides voltage waveforms, making it suitable for frequency analysis in a manner analogous to conventional MEA recordings. Figure 4A displays a representative raw waveform recorded from a single electrode in a 5-month-old cerebral organoid, alongside corresponding signals filtered into delta (0.5–3 Hz), theta (4–7 Hz), alpha (8– 11 Hz), beta (12–29 Hz), and gamma (30–100 Hz) bands using a finite impulse response (FIR) bandpass filter. When a network burst occurred across the entire cortical organoid, slow-wave field potentials were observed, as shown in Figure 4A. Field potentials were acquired from 46,630 electrodes, enabling frequency analysis at the whole-organoid level (Figure 4B). Figure 4B shows heatmaps of standardized potential intensities for each frequency band across all electrodes. Delta to beta bands exhibited a gradient of increasing intensity from the outer edge toward the center, with the strongest signals observed in the lower-right region. In contrast, gamma-band activity showed a distinct pattern, characterized by weaker signals in the lower half and stronger signals in the upper half of the organoid. To delineate frequency-specific spatial domains within the organoid, k-means clustering analysis was performed, segmenting the organoid into nine distinct clusters (Figure. 4C). Comparison of the frequency intensity profiles across clusters revealed that Clusters 1–3 exhibited low intensity across all bands, whereas Clusters 5, 7, and 8 showed uniformly high intensity. Clusters 6 and 9 exhibited high intensity in the delta to beta bands but relatively low gamma-band activity (Figure. 4D). Frequency analysis of activity waveforms detected from 46,630 electrodes on the UHD-CMOS MEA demonstrated that region-specific frequency characteristics can be extracted from the brain organoid.

**Figure 4.**
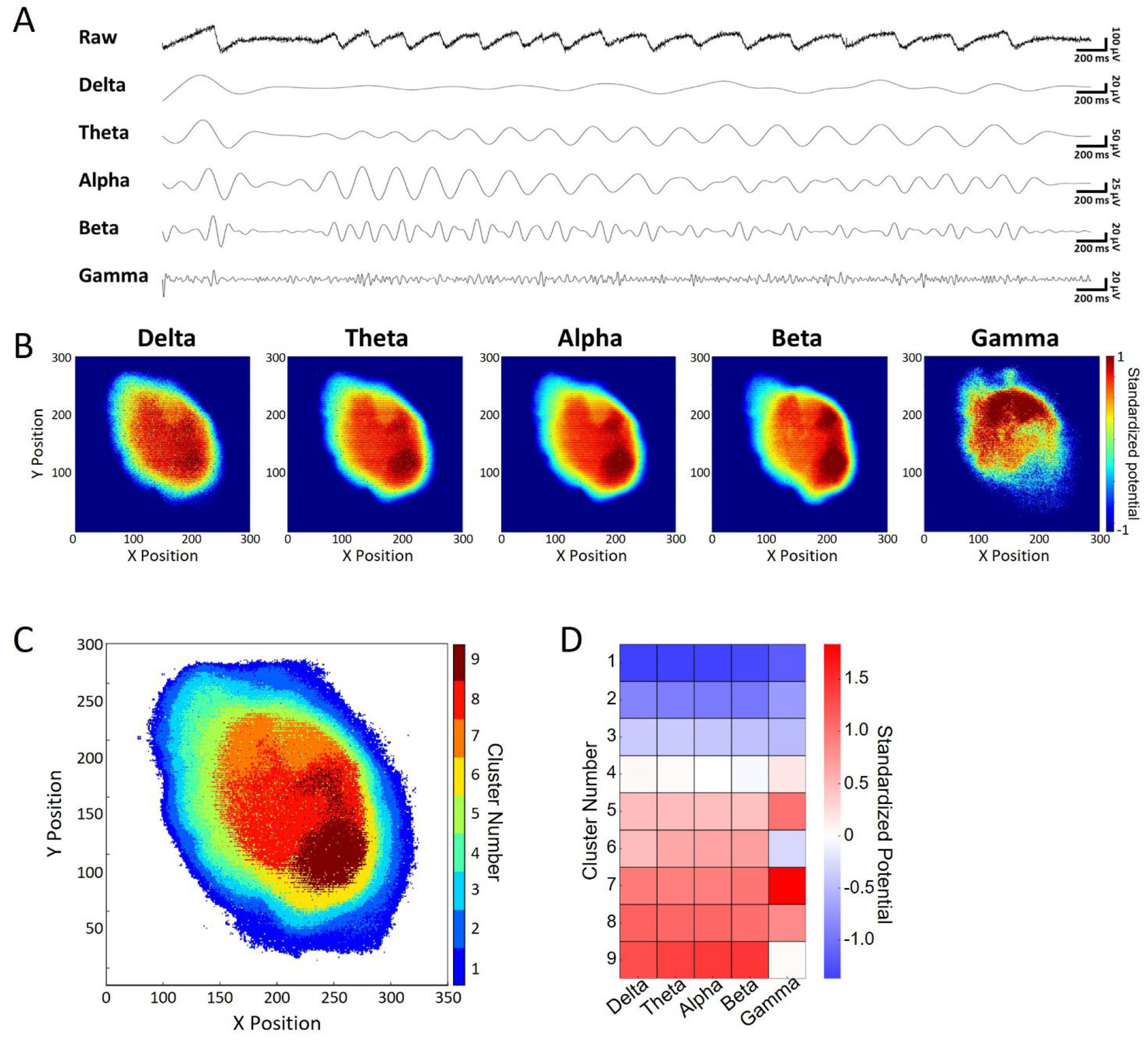
Frequency distribution characteristics of cortical organoids. (A) Spontaneous oscillation waveforms for each frequency band. From top to bottom: raw waveform, delta, theta, alpha, beta, gamma. (B) Spatial distribution maps of normalized signal intensity for each frequency band. (C) Clustering map showing nine groups classified based on the intensity profiles of five frequency bands. (D) Heatmap of frequency band intensities for each cluster.

### 3.4 Inter-Regional Connection Analysis in Midbrain–Striatal Assembloids

Using the UHD-CMOS MEA, we recorded the electrical activity of midbrain–striatal assembloids and evaluated inter-regional connection strength based on single-cell firing activity. Figure 5A shows a heatmap of peak voltage distribution over a one-minute recording period, where the upper part corresponds to the striatum and the lower part to the midbrain organoid. Electrical activity was detected in both regions, indicating that each tissue was functionally active. To assess the effects of pharmacological intervention, 30 µM of 4-aminopyridine (4-AP), a potassium channel blocker, was administered to the assembloid, and changes in firing frequency and inter-regional connectivity were evaluated. Figure 5B presents raster plots of individual cells (red: striatum, yellow: midbrain). Following 4-AP administration, spike counts increased in both the midbrain and striatum (Figure. 5C). Connection strength was calculated by normalizing the number of synchronized spikes (defined as spikes occurring within 100 ms between neuron pairs) by the average firing rate of each pair. Figure 5D shows the connection strength within the striatum, within the midbrain, and between the midbrain and striatum before and after 4-AP treatment. The connection strength increased from 0.0360 to 0.0946 within the striatum and from 0.181 to 0.377 within the midbrain. Furthermore, the inter-regional connection strength between the midbrain and striatum also increased from 0.0305 to 0.0632. Single-cell-based network connectivity analysis using the UHD-CMOS MEA revealed that changes in inter-organoid information transmission within assembloids can be detected following compound administration.

**Figure 5.**
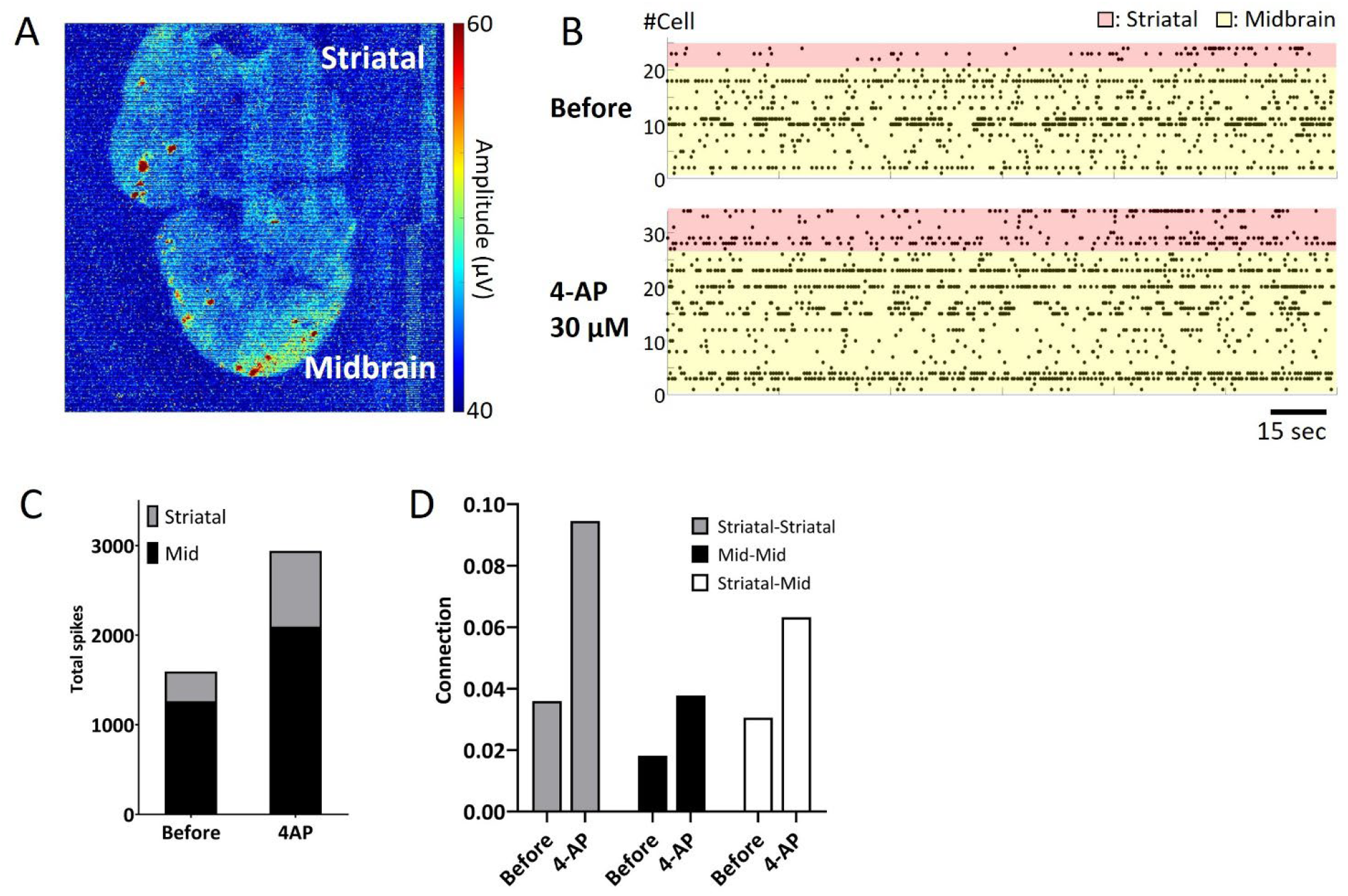
Analysis of inter-regional connectivity in midbrain–striatal assembloids. (A) Peak potential map of spontaneous activity over a 1-minute recording. Upper region: striatum; lower region: midbrain. (B) Raster plot of spontaneous spikes from individual cells over 1 minute. Red: striatum; yellow: midbrain. Scale bar = 15 sec. (C) Total spike count before and after 4-AP administration. The total number of spikes per minute is shown for each region. Gray: striatum; black: midbrain. (D) Connection strength before and after 4-AP administration. Strength was calculated by normalizing the number of synchronous spikes (within 100 ms) by the mean spike count of each cell pair. Gray: within striatum; black: within midbrain; white: between midbrain and striatum.

## 4 Discussion

In this study, we performed large-scale recordings of brain organoids using field potential imaging (FPI) with an ultra-high-density (UHD) CMOS microelectrode array (MEA) comprising 236,880 microelectrodes (each measuring 10.52 μm × 10.52 μm) and covering a broad sensing area of 32.45 mm^2^, in order to precisely characterize the detailed electrical activity of brain organoids. Based on single-cell activity, we analyzed neuronal network connectivity, propagation velocity, propagation area, and frequency characteristics.

L-Dopa, a precursor to dopamine, is taken up by neurons and decarboxylated by Aromatic L-Amino Acid Decarboxylase (AADC), resulting in the production of dopamine, which is subsequently released into the extracellular space. When L-Dopa was administered to midbrain organoids, a decrease in activity was observed in 10.1% of cells at a concentration of 0.3 µM (Figure 1E). TH-positive dopaminergic neurons are known to express D2 autoreceptors (Beaulieu and Gainetdinov, 2011), and this suppression of activity is thought to be due to D2 receptor activation following dopamine release. However, at higher concentrations, this suppression was no longer observed (Figure. 1E), and instead, an enhancement of network activity was seen throughout the midbrain organoids (Figure. 2A). The midbrain organoids used in this study contained not only TH-positive neurons (Figure. 1A) but also likely included vglut2-positive glutamatergic neurons, which are known to be present in the midbrain, as well as cells expressing D1-type dopamine receptors (Yamaguchi et al., 2007; Yamaguchi et al., 2011; Morales and Root, 2014). It is possible that excitatory input from these cells indirectly disinhibited or activated dopaminergic neurons, leading to increased activity at the single-cell level (Figure. 1E) and strengthened network connectivity across the entire organoid (Figure. 2A). The use of UHD CMOS MEA enabled quantitative evaluation of both the firing characteristics of individual cells and the neural network activity based on single-cell firing patterns within the brain organoids.

Based on the capabilities of the UHD-CMOS-MEA, novel endpoints for the propagation velocity and area of network activity were established. In cortical organoids, the propagation velocity of network activity increased following administration of picrotoxin (PTX), a GABA_A_ receptor antagonist. This observation aligns with previous reports showing that administration of bicuculline methiodide— another GABA_A_ receptor antagonist—increased propagation velocity in cortical slices derived from human epilepsy patients (Covelo et al., 2025). Although the compounds differ, the results may reflect similar physiological effects mediated by GABA_A_ receptor inhibition. Following administration of MK-801, an NMDA receptor antagonist, a reduction in the propagation area was observed in most regions of the organoids (Figure 3D, E). Similar reductions in activity propagation caused by NMDA receptor antagonists have been reported in in vivo EEG recordings from rats and in brain slice experiments (Morales-Villagrán et al., 1996; Fukuda et al., 1998), suggesting that the responses observed in this study may reflect physiological phenomena in the intact brain. Notably, although the propagation area decreased across the majority of the organoid, a localized enhancement in propagation was detected in the lower-right region following treatment with 1 µM MK-801 (Figure 3D). This focal increase in activity may be attributable to disinhibition resulting from impaired function of GABAergic neurons. Previous studies have described both direct suppression of inhibitory neuron activity via NMDA receptor blockade and indirect effects through reduced excitatory input (Li et al., 2002; Homayoun and Moghaddam, 2007). These mechanisms are considered to underlie the excitotoxic effects of NMDA receptor antagonists. Due to self-organization, brain organoids likely exhibit regional variability in the ratio of excitatory to inhibitory neurons. Consequently, in areas with a lower proportion of inhibitory neurons, the loss of inhibitory control induced by MK-801 may be more pronounced, potentially leading to enhanced excitatory activity mediated by AMPA receptors. Analysis of propagation velocity and propagation area using the UHD-CMOS MEA demonstrated that this method is effective for evaluating compound responses related to synaptic transmission in brain organoids. Moving forward, integrating analyses of propagation pathways may enable the identification of regions exhibiting disease-related activity abnormalities or neural circuits serving as initiation sites for compound responses.

Since FPI represents voltage waveforms, frequency analysis can be conducted similarly to conventional MEA measurements. Traditional frequency analyses of brain organoids using standard MEAs were limited by the small number of electrodes, typically ranging from 16 to 64. In this study, activity was acquired from 46,630 electrodes using the UHD CMOS MEA, which enabled a comprehensive analysis of frequency distributions across the entire organoid. Delta band potentials increased from the periphery toward the center. Alpha to beta band activity also intensified toward the center, with particularly strong signals observed in the lower-right region of the organoid. In contrast, gamma band activity displayed a distinct pattern—being stronger in the upper region and weaker in the lower region (Figure 4B). Clustering analysis enabled visualization of the organoid’s structural features based on frequency-specific characteristics (Figure 4C). However, because the organoids were measured without being anchored to the electrodes, the variability in adhesion between the organoid and electrodes must be considered. The periphery of the organoid likely had fewer cells adhering to the electrodes, which may explain the consistently weak potentials observed in clusters 1, 2, and 3 across all frequency bands. In contrast, clusters 6, 7, 8, and 9, located deeper within the organoid, exhibited variations in gamma band potentials (Figure 4C), indicating that in regions with better electrode contact, the intrinsic structural frequency properties of the organoid were more accurately reflected. The distribution of gamma band activity may be associated with the expression of PV-positive GABAergic neurons (Cardin et al., 2009), which are known to be essential for gamma wave generation in the cortex. In clusters 6 and 9, where gamma activity was weak while other frequency bands were strong, it is likely that PV-positive GABAergic neurons were either absent or functionally immature. Previous studies have shown that the expression of GABAergic neurons in cortical organoids increases after six months of culture (Trujillo et al., 2019). Given that the organoids in this study were five months old, it is plausible that some regions lacked sufficient expression or functional maturity of PV-positive GABAergic neurons. Comparing the electrophysiological frequency characteristics of organoids with cell type identity and the histochemically organized circuit architecture within the organoids remains an important challenge for future studies. Moreover, samples that exhibited global oscillatory activity throughout the entire organoid, as observed here, were extremely rare. Even when derived from the same iPSC dish, some organoids exhibited oscillations only in localized areas, while others showed no detectable electrical activity. Therefore, establishing reproducible methods for reliably generating organoids with consistent, whole-organoid activity remains a critical goal. Despite these challenges, this study demonstrates that UHD CMOS MEA technology enables comprehensive frequency distribution analysis of entire brain organoids. Importantly, the ability to detect spatial characteristics of gamma band activity—crucial for the evaluation of neurological disorders such as epilepsy, Alzheimer’s disease, and autism—highlights the utility of this method for assessing frequency-specific disease models (Herrmann and Demiralp, 2005; Başar, 2013; McNally and McCarley, 2016; Guan et al., 2022).

Currently, the evaluation of assembloids is primarily performed using techniques such as optogenetic observation with adeno-associated viruses (AAV), calcium imaging, patch-clamp recordings, and conventional multi-electrode arrays (MEAs) (Andersen et al., 2020; Miura et al., 2020; Reumann et al., 2023; Ozgun et al., 2024). However, these methods have limitations in terms of temporal and spatial resolution, as well as invasiveness to the cells. In contrast, the UHD CMOS MEA enabled the non-invasive detection of electrical activity at single-cell resolution across the entire assembloid, with both high temporal and spatial precision (Figure 5A). When 4-Aminopyridine (4-AP), a potassium channel blocker, was applied to midbrain–striatal assembloids, increased firing activity was observed in both the midbrain and striatal regions (Figure 5B). Furthermore, an enhancement of inter-regional connectivity was also detected (Figure 5C). A current limitation of this experimental setup is the inability to selectively apply compounds to individual tissues, leading to simultaneous exposure of both regions. This complicates the mechanistic analysis of inter-regional connection enhancement. One potential solution is to combine UHD CMOS MEA recordings with optogenetic techniques, such as AAV-mediated gene delivery, enabling targeted stimulation of specific cell types and subsequent recording of their responses. Additionally, integrating UHD CMOS MEA with Microphysiological System (MPS) devices containing microfluidic channels would allow for region-specific compound administration. MPS platforms seed organoids into individual chambers connected via microfluidic pathways, offering both structural control and the ability to measure not only tissue-specific activity but also axonal signals transmitted through the connecting channels. A system that combines these technologies with UHD CMOS MEA for assessing inter-organ connectivity in assembloids may be particularly valuable for investigating projection deficits caused by neurological disorders. It may also be applied to disease models involving peripheral nervous system damage, such as those using sensory neuron–spinal cord assembloids.

In this study, we established a methodology for large-scale recording of brain organoids using UHD-CMOS MEA, along with analytical techniques to assess network connectivity, propagation velocity, propagation area, and frequency dynamics at the single-cell level. While further refinement is needed to achieve stable organoid generation and recording techniques, and to elucidate the relationship between electrophysiological activity and structural characteristics, this approach holds promise for advancing our understanding of the functional electrophysiology of brain organoids and assembloids. Moreover, it is expected to be valuable for compound screening and the evaluation of human neurological disease models.

## 5 Conflict of Interest

I.S. is a founder of VitroVo, Inc. R.Y., N.M. and Y.I. are also employees of VitoroVo. Inc.

## 6 Author Contributions

Conceptualization, R.Y. and I.S.; writing—original draft preparation, R.Y. and I.S.; writing—review and editing R.Y., I.S., Y.I. and N.M.; Formal analysis, R.Y., Y.I. and N.M.; cell line culture, R.Y.; immunostaining, R.Y.; funding acquisition, I.S. All authors have read and agreed to the published version of the manuscript.

## 7 Funding

This study was supported by the Japan Society for the Promotion of Science (JSPS) KAKENHI (Grant Number 23H03726 and 25K21561) and collaboration research grant with Sony semiconductor solutions Inc.

## 8 Acknowledgments

This study was supported by the Japan Society for the Promotion of Science (JSPS) KAKENHI Grant Number 23H03726 and 25K21561. This study was also supported by the UHD-CMOS-MEA system with Sony semiconductor solutions Inc.

## 9 Data Availability Statement

The data and scripts that support the findings of this study are available from the corresponding author upon reasonable request.

## Notes

### Summary of Updates

We have revised the manuscript to enhance clarity and consistency. Minor textual edits have been made throughout the main text. Additionally, the Competing Interests, Funding, and Acknowledgments sections have been updated to reflect accurate and current information.

