## Supplementary Material for "Advanced neural activity mapping in brain organoids via field potential imaging with ultra-high-density CMOS microelectrodes"

**Supplementary Table 1.** Composition of culture media used for cerebral organoid generation, prepared according to the STEMdiff™ Cerebral Organoid Kit protocol (08570, STEMCELL Technologies), including Y-27632 as specified in the manufacturer's instructions.

| **Medium** | **Component** | **Source** | **Cat. No.** | **Amount** |
| --- | --- | --- | --- | --- |
| EB seeding medium | STEMdiff Cerebral Organoid Supplement A | STEMCELL Technologies | 08574 | 10 mL |
|  | STEMdiff Cerebral Organoid Basal Medium 1 | STEMCELL Technologies | 08572 | 40 mL |
|  | Y-27632 (10mM) | FUJIFILM Wako | 036-24023 | 0.05 mL |
| EB Formation Medium | STEMdiff Cerebral Organoid Supplement A | STEMCELL Technologies | 08574 | 10 mL |
|  | STEMdiff Cerebral Organoid Basal Medium 1 | STEMCELL Technologies | 08572 | 40 mL |
| Induction Medium | STEMdiff Cerebral Organoid Supplement B | STEMCELL Technologies | 08575 | 0.5 mL |
|  | STEMdiff Cerebral Organoid Basal Medium 1 | STEMCELL Technologies | 08572 | 49.5 mL |
| Expansion Medium | STEMdiff Cerebral Organoid Supplement C | STEMCELL Technologies | 08576 | 0.25 mL |
|  | STEMdiff Cerebral Organoid Supplement D | STEMCELL Technologies | 08577 | 0.5 mL |
|  | STEMdiff Cerebral Organoid Basal Medium 2 | STEMCELL Technologies | 08573 | 24.25 mL |
| Maturation Medium | STEMdiff Cerebral Organoid Supplement E | STEMCELL Technologies | 08578 | 4.5 mL |
|  | STEMdiff Cerebral Organoid Basal Medium 2 | STEMCELL Technologies | 08573 | 224.5 mL |

**Supplementary Table 2.** Composition of culture media for midbrain–striatal assembloid generation using STEMdiff™ organoid kits (100-1096 and 08620, STEMCELL Technologies), including Y-27632 and region-specific supplements recommended by the manufacturer.

| **Medium** | **Component** | **Source** | **Cat. No.** | **Amount** |
| --- | --- | --- | --- | --- |
| Organoid Formation Medium | STEMdiff™ Neural Organoid Basal Medium 1 | STEMCELL Technologies | 08621 | 20 mL |
| Midbrain Organoid Expansion Medium | STEMdiff™ Neural Organoid Basal Medium 2 | STEMCELL Technologies | 08622 | 49 mL |
|  | STEMdiff™ Neural Organoid Supplement A | STEMCELL Technologies | 08623 | 1 mL |
|  | STEMdiff™ Neural Organoid Supplement K | STEMCELL Technologies | 100-1094 | 0.1 mL |
|  | STEMdiff™ Neural Organoid Supplement L | STEMCELL Technologies | 100-1095 | 0.1 mL |
| Striatal Organoid Expansion Medium | STEMdiff™ Neural Organoid Basal Medium 2 | STEMCELL Technologies | 08622 | 49 mL |
|  | STEMdiff™ Neural Organoid Supplement A | STEMCELL Technologies | 08623 | 1 mL |
|  | Actin A (10 µg/ml) | PeproTech | 120-14P | 0.25 mL |
|  | IWP-2 (2.5 mM) | Selleck Chemicals | S7085 | 0.05 mL |
|  | SR11237 (100 µM) | Sigma Aldrich | S8951 | 0.05 mL |
| Organoid Differentiation Medium | STEMdiff™ Neural Organoid Basal Medium 2 | STEMCELL Technologies | 08622 | 49 mL |
|  | STEMdiff™ Neural Organoid Supplement A | STEMCELL Technologies | 08623 | 1 mL |
|  | STEMdiff™ Neural Organoid Supplement C | STEMCELL Technologies | 08625 | 0.05 mL |
| Organoid Maintenance Medium | STEMdiff™ Neural Organoid Basal Medium 2 | STEMCELL Technologies | 08622 | 49 mL |
|  | STEMdiff™ Neural Organoid Supplement A | STEMCELL Technologies | 08623 | 1 mL |
